# Understanding inhibitor resistance in Mps1 kinase through novel biophysical assays and structures

**DOI:** 10.1101/112086

**Authors:** Yoshitaka Hiruma, Andre Koch, Nazila Hazraty, Foteini Tsakou, René H. Medema, Robbie P. Joosten, Anastassis Perrakis

## Abstract

Monopolar spindle 1 (Mps1/TTK) is a protein kinase essential in mitotic checkpoint signalling, preventing anaphase until all chromosomes are properly attached to spindle microtubules. Mps1 has emerged as a potential target for cancer therapy, and a variety of compounds have been developed to inhibit its kinase activity. Mutations in the catalytic domain of Mps1 that give rise to inhibitor resistance, but retain catalytic activity and do not display cross-resistance to other Mps1 inhibitors, have been described. Here we characterize the interactions of two such mutants, Mps1 C604Y and C604W, which raise resistance to two closely related compounds, NMS-P715 and its derivative Cpd-5, but not to the well-characterised Mps1 inhibitor, reversine. We show that estimates of the IC_50_ (employing a novel specific and efficient assay that utilizes a fluorescently labelled substrate) and of the binding affinity (*K*_D_) indicate that in both mutants, Cpd-5 should be better tolerated than the closely related NMS-P715. To gain further insight, we determined the crystal structure of the Mps1 kinase mutants bound to Cpd-5 and NMS-P715, and compare the binding modes of Cpd-5, NMS-P715 and reversine. The difference in steric hindrance between Tyr/Trp604 and the trifluoromethoxy moiety of NMS-P715, the methoxy moiety of Cpd-5, and complete absence of such a group in reversine, account for differences we observe *in vitro*. Our analysis enforces the notion that inhibitors targeting Mps1 drug-resistant mutations can emerge as a feasible intervention strategy based on existing scaffolds, if the clinical need arises.

**Summary statement:** The inhibition of specific Mps1 kinase inhibitors towards the wild-type protein and inhibitor-resistant mutants is explained by a novel specific activity assay, biophysical characterisation, and X-ray structures.

**Abbreviations:**

ATP: adenosine triphosphate
Bub1/Bub3: budding uninhibited by benzimidazoles 1 / budding uninhibited by benzimidazoles 3
Cpd-5: Compound-5 (N-(2,6-diethylphenyl)-8-((2-methoxy-4-(4-methylpiperazin-1-yl)phenyl)amino)-1-methyl-4,5-dihydro-1H-pyrazolo[4,3-h]quinazoline-3-carboxamide)
FP: fluorescence polarization
GST: Glutathione S-transferase
IC50: half maximal inhibitory concentration
K_D_: dissociation constant
ΔG_calc_: calculated Gibbs energy difference for ligand binding
KNL1: kinetochore null protein 1
KPi: (inorganic) potassium phosphate
Mps1: monopolar spindle 1
MST: microscale thermophoresis
NMS-P715: N-(2,6-diethylphenyl)-1-methyl-8-({4-[(1-methylpiperidin-4-yl)carbamoyl]-2-(trifluoromethoxy)phenyl}amino)-4,5-dihydro-1H-pyrazolo[4,3-h]quinazoline-3-carboxamide
TMR: tetramethylrhodamine
WT: wild-type

## Introduction

One of the common features of solid human tumors is presence of abnormal number of chromosomes, a state often referred as “aneuploidy” [1]. Previous studies indicate that a mechanism that sustains aneuploidy in tumour cells is the over-expression of high levels of monopolar spindle 1 (Mps1) kinase [2]. Mps1, also known as threonine and tyrosine kinase (TTK), is a dual specificity protein kinase, essential for safeguarding proper chromosome alignment and segregation during mitosis [2]. Due to its importance for the viability of tumor cells, Mps1 kinase has become a potential target for cancer treatment [2]. Over the past years, several dozens of small compounds have been developed to inhibit the Mps1 kinase activity [2–7]. Recently, NMS-P715 has been described to suppress the growth of medulloblastoma cells, a common malignant brain tumor in children [8]. The anti-proliferative activity of NMS-P715 has been also shown in breast, renal and colon cancer cell lines [9]. A derivative of NMS-P715, Compound 5 (Cpd-5) [10], has been reported to display higher potency towards Mps1 than NMS-P715 [11]. Although these Mps1 inhibitors have promising results in pre-clinical studies [4,8,9], they also have innate problems as kinase inhibitors [11,12]. After the initial drug response, tumor cells frequently acquire resistance and become insensitive to treatment [11,13]. The development of drug resistance in cancer cells is often the consequence of mutations of the targeted kinases [14]. These mutations are typically found in the ATP binding pocket, which renders the binding of inhibitors suboptimal, while retaining kinase activity [15]. A mutation at Cys604 of the Mps1 kinase has been isolated by raising resistance against a number of inhibitors including NMS-P715 [12] as well as Cpd-5 [11], in cell studies. It is positioned in the “hinge loop” region of the kinase domain, which is part of the ATP binding pocket [2]. Gurden *et al.* described the isolation of the C604W mutant [12], and a C604Y mutation has been independently described by Koch *et al.*, raising resistance to Cpd-5 [11]. A crystal structure of the Mps1 kinase C604W mutant bound to NMS-P715 [12] showed how this mutation prohibits efficient binding of NMS-P715, suggesting that further development of NMS-P715 derivatives could prove necessary to combat tumor cells conferring drug resistances.

Drug development against kinases can be made more efficient by the availability of automatable direct-readout specific assays, which can be implemented in common laboratory equipment. To detect Mps1 kinase activity *in vitro*, the myelin basic protein (MBP) is widely employed as a substrate, using radiolabelled ATP or specific phospho-Mps1 antibodies [16–18]. Alternatively, the KNL1 protein is used as substrate [11], which is a well-documented Mps1 substrate of kinetochore components [19,20]. These assays are highly specific for Mps1, but hard to quantify. Another more recently demonstrated way of measuring Mps1 kinase activity is a mobility shift assay described by Naud *et al* [5]: phosphorylated and non-phosphorylated peptides are separated by electrophoresis based on their charges [14]. This mobility shift assay has been compared with a radiometric assay and found to be more robust [21].

Here, we first describe a novel and highly specific assay for Mps1 activity by using fluorescence polarization (FP), which is based on the change of tumbling rate of a fluorescently-labelled KNL1 peptide, monitoring the binding of the 73 kDa Bub1/Bub3 complex as a high-throughput highly specific physiological readout. Using this assay, we determined the IC_50_ of NMS-P715 and Cpd-5 for the Mps1 WT and C604Y/W mutant, and compare it to the *K*_D_ against these Mps1 variants. We also present the crystal structures of the Mps1 kinase domain variants bound to Cpd-5 and NMS-P715 and discuss the structural basis of the inhibition mode of Cpd-5 over NMS-P715 and reversine in the Mps1 C604Y/W mutants, and how differences between the binding modes of related inhibitors could be exploited in therapy.

## Materials and Methods

### Protein production

The plasmid containing a construct of the Mps1 kinase domain (residues 519–808) was a gift from Dr. Nicola Burgess-Brown (Addgene plasmid # 38907) [22]. The site-specific mutant of C604W was generated using the QuickChange (Stratagene). The Mps1 kinase domain C604Y/W variants were produced as previously described [23].

The pFastBac plasmid containing the GST-tagged full-length Mps1 was a gift from Dr. Geert Kops (Hubrecht Institute, Utrecht, the Netherlands). Recombinant baculovirus was produced according to the Bac-to-Bac protocols (Invitrogen). *Spodoptera frugiperda* (Sf9) insect cells were infected with the baculovirus and allowed to grow for 72 hours at 27 °C. Cells were harvested by centrifugation and re-suspended in 50 mL of 20 mM KP_i_, pH 7.5, 150 mM KCl and 1 mM TCEP (buffer A) supplemented with one tablet of Pierce™ Protease Inhibitor Tablets EDTA-free (Thermo Fisher Scientific). Samples were stored at –20°C before proceeding to purification. The re-suspended cells were defrosted at room temperature and lysed by sonication for 1 min at 50% amplitude in a Qsonica Sonicator Q700 (Fisher Scientific). Following centrifugations at 21000 *g* for 20 minutes at 4 °C, the supernatant was loaded on Glutathione Sepharose 4B (GE Healthcare). After extensive washing in buffer A, the protein was eluted in buffer A supplemented with 10 mM GSH. The sample was then loaded on a Superdex G75 16/60 HiLoad (GE Healthcare) pre-equilibrated in 20 mM HEPES/NaOH, 50 mM KCl, and 3 mM DTT. The protein fractions were pooled together and concentrated to ~10 μM. The concentration of the GST-Mps1 full length was determined by absorption spectrophotometry at 280 nm, with calculated ε = 122.8 mM^-1^ cm^-1^. The purified protein was aliquoted and stored at -–80°C.

Bub1 (residues 1–280) and Bub3 were produced as previously described [19], with modifications. The baculovirus with the Bub1 and Bub3 constructs was a gift from Dr. Geert Kops. Sf9 insect cells were infected and allowed to grow for 72 hours at 27 °C. The cell cultures were harvested by centrifugation and re-suspended in 50 mM Tris/HCl, pH 7.7, 1 mM TCEP and 0.05% Tween20 (buffer B) supplemented with 300 mM KCl, 10 mM imidazole and one tablet of Pierce™ Protease Inhibitor Tablets EDTA-free (Thermo Fisher Scientific). Samples were stored at –20°C before proceeding to the purification. The re-suspended cells were defrosted at room temperature and lysed by sonication for 1 min at 50% amplitude in a Qsonica Sonicator Q700 (Fisher Scientific). Following centrifugation at 21000 *g* for 20 minutes at 4 °C, the supernatant was loaded on a HisTrap HP column (GE Healthcare). After extensive washing in buffer B supplemented with 300 mM KCl and 10 mM imidazole, the protein was eluted with buffer B supplemented with 150 mM imidazole. The sample was then loaded on a Superdex G75 16/60 HiLoad (GE Healthcare) pre-equilibrated in a buffer B supplemented with 100 mM KCl. The protein fractions were pooled together and concentrated. The concentration of the Bub1/Bub3 complex was determined by absorption spectrophotometry at 280 nm, with calculated ε = 101.9 mM^-1^ cm^-1^. The purified protein was aliquoted and stored at –80°C.

### Chemicals

Cpd-5 was synthesized as previously described [10]. NMS-P715 and reversine were purchased from Merck Millipore and Sigma-Aldrich, respectively. A KNL1 peptide consisting of 31 residues (^162^KHANDQTVIFSDENQMDLTSSHTVMITKGLK^191^) was chemically synthesized and labelled with Tetramethylrhodamine (TMR) at the N-terminus (TMR-KNL1_p_).

### Western4blot based Mps1 kinase activity assay

100 ng of Mps1 wild type or Mps1 C604Y was incubated for 10 min at 32°C with the indicated compounds in a buffer containing 50 mM Tris-HCl pH 7.5, 100 mM NaCl, 20 mM MgCl_2_, 1 mM DTT, 0.2 mM ATP. After this incubation, 200 ng of purified GST-KNL1-M3 was added, and the mixture was incubated for 60 min at 32°C before addition of SDS-sample buffer and heating at 95°C for 10 min. Samples were separated by SDS-PAGE and transferred to nitrocellulose membranes, blocked with 4% (w/v) bovine serum albumin at room temperature for 0.5 hour, and incubated with primary antibody (phospho-KNL1 MELT13/17; a kind gift from Dr. Geert Kops [24] at 4°C overnight. After incubation with secondary antibody (1:2000 dilution) at room temperature for 1 hour, the membranes were developed with chemiluminescence ECL reagent (Amersham) and pictures were taken with the ChemiDOC XRS+ (BioRad). Resulting images were analysed using ImageJ software.

### Fluorescence-based Mps1 kinase activity assay

The Bub1/Bub3 complex was titrated to a reaction mixture containing 20 nM TMR-KNL1_p_ and 10 nM GST-Mps1 full length variants in 20 mM HEPES/NaOH pH 7.4, 1 mM ATP, 4 mM MgCl_2_, 200 μM TCEP and 0.05% Tween20 (buffer C) with or without 50 nM of inhibitors. The samples were incubated at room temperature in a 384 well Corning assay plate. All measurements were performed in a Pherastar plate reader (BMG LABTECH GmbH, Germany).The excitation and emission wavelengths were 540 nm and 590 nm, respectively and fluorescence polarization (FP) was calculated. All measurements were carried out in duplicate.

To calculate the IC_50_ values, the inhibitors were titrated to a reaction mixture containing 20 nM TMR-KNL1 peptide, 200 *n*M Bub1/Bub3 complex and 5 nM GST-Mps1 full length variants in buffer C. The samples were incubated at room temperature overnight. Measurements were carried in duplicates. The FP values were plotted against inhibitor concentration and fitted with a standard one site model (equation 1) using non-linear regression in GraphPad Prism 6 (GraphPad Software, Inc, USA).

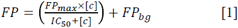

where FP_max_ is maximum binding in FP; [c] is inhibitor concentration; FP_bg_ is the background FP value.

### Microscale Thermophoresis

The thermophoresis measurements and data analysis were performed as previously described [23] with a slight modification. The DY-547P1 labelled samples were used at a final concentration of 20 nM in Tris buffer (50 mM Tris/HCl, pH 7.4, 150 mM KCl, 1% DMSO, 0.05% Tween20). The measurement was performed at 20% LED and 40% MST power.

### Crystallization

The purified Mps1 kinase domain C604Y/W variants were co-crystallized with Cpd-5 as well as NMS-P715, using the sitting drop vapour diffusion method in MRC 2-Well Crystallization Plate (Swissci) UVP plates (Hampton Research), with standard screening procedures [25]. The protein solution at ~200 μM (~7.2 mg mL^-1^) was pre-incubated with 250 μM of Cpd-5 or NMSP715, and 0.1 μL of this solution was mixed with 0.1 μL of reservoir solution and equilibrated against a 50 μL reservoir. Crystals were obtained in 10–17% (w/v) PEG 350 MME, 1–10 mM MgCl_2_, and 100 mM Tris/HCl, pH 7.5–9. Crystals appeared at 18 °C within 72 hours. Crystals were briefly transferred to a cryoprotectant solution containing the reservoir solution and 20% (w/v) ethylene glycol and vitrified by dipping in liquid nitrogen.

### Data collection and structure refinement

X-ray data were collected on beamline ID30A-3, ID29 and ID30A-1 at the European Synchrotron Radiation Facility (ESRF) for 5MRB, 5NTT and 5O91 (PDB ID codes) respectively. The images were integrated with XDS [26] and merged and scaled with AIMLESS [27]. The starting phases were obtained by molecular replacement using PHASER [28] with an available Mps1 structure (PDB, 3HMN) [29] as the search model. Geometric restraints for the compounds were made in AceDRG [30]. The models were built using COOT [31] and refined with REFMAC [32] in iterative cycles. Model re-building and refinement parameter adjustment were performed in PDB-REDO [33,34]; homology-based hydrogen bond restraints (van Beusekom et al; DOI: 10.1101/147231) were used at some stages of the procedure. The quality of the models was evaluated by MOLPROBITY [35]. Data collection and refinement statistics are presented in Table 1.

**Table 1.** Crystallographic Data. High Resolution shell in parentheses

| Protein complex | Cpd5-C604Y | NMS-C604Y | Cpd5-C604W |
| --- | --- | --- | --- |
| PDB identifier | 5MRB | 5NTT | 5O91 |
| Data Collection |  |  |  |
| Wavelength (Å) | 0.97 | 1.07 | 0.97 |
| Resolution (Å) | 41.39–2.20<br>(2.27–2.20) | 41.65–2.75<br>(2.90–2.75) | 41.76–3.20<br>(3.42–3.20) |
| Space Group | I 2 2 2 | I 2 2 2 | I 2 2 2 |
| Unit Cell a, b, c (Å) | 70.58, 111.42,<br>115.03 | 70.67, 112.01,<br>116.13 | 70.67, 111.89,<br>116.80 |
| CC<sub>1/2</sub> | 0.999 (0.702) | 0.996 (0.571) | 0.985 (0.538) |
| R<sub>merge</sub> | 0.040 (0.764) | 0.138 (1.283) | 0.258 (1.595) |
| I/σI | 18.2 (1.9) | 9.6 (1.2) | 6.0 (1.0) |
| Completeness (%) | 99.3 (99.8) | 100 (100) | 100 (100) |
| Multiplicity | 4.6 (4.8) | 5.9 (5.8) | 6.4 (6.3) |
| Unique Reflections | 23225 | 12326 | 7947 |
| Refinement |  |  |  |
| Atoms<br>protein/ligand/other | 2164/43/119 | 2278/49/40 | 2185/43/14 |
| B-factors<br>protein/ligand/other (Å<sup>2</sup>) | 62/53/69 | 76/88/60 | 81/95/44 |
| R<sub>work</sub>/R<sub>free</sub> (%) | 17.3/21.6 | 19.9/23.0 | 21.2/25.4 |
| Bonds rmsZ/rmsd (Å) | 0.525/0.011 | 0.381/0.008 | 0.381/0.008 |
| Angles rmsZ/rmsd (°) | 0.637/1.366 | 0.563/1.235 | 0.547/1.189 |
| Ramachandran<br>preferred/outliers (%) | 97.7/0.4 | 98.5/0.4 | 96.2/0.0 |
| Rotamers<br>preferred/outliers (%) | 96.7/0.8 | 96.5/0.8 | 91.9/1.2 |
| Clash score (%-ile) | 100 | 100 | 100 |
| MolProbity score (%-ile) | 100 | 100 | 100 |

## Results and Discussion

### Cpd-5 and NMS-P715 inhibit Mps1-mediated phosphorylation of KNL1 peptides

To enable quick spectroscopic quantitation of Mps1 activity, we synthesized a fluorescent KNL1 peptide, TMR-KNL1. As shown in Figure 1A, phosphorylation of TMR-KNL1 by Mps1 can be detected specifically by the Bub1/Bub3 complex that binds only the phosphorylated form of the peptide. Bub1/Bub3 form a tight complex with TMR-KNL1, with a dissociation constant (*K*_D_) of 12.2±1.4 nM (Figure 1B). Binding of the 73 kD Bub1/Bub3 complex reduces the tumbling rate of the ~4 kD TMR-KNL1 peptide, allowing measurement of Bub1/Bub3 binding, and therefore of phosphorylation, by fluorescence polarization (FP). Addition of the Cpd-5 or NMS-P715 Mps1 inhibitors does not notably change the binding constant of the Bub1/Bub3 complex, as expected, but only the end point measurement of the FP signal, which is proportional to the degree of phosphorylation of the peptide.

**Figure 1:**
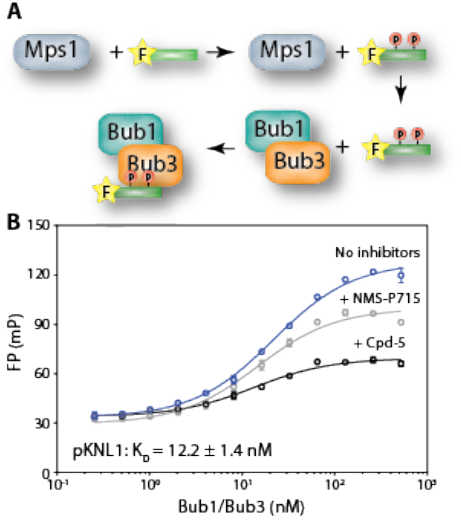
fluorescence polarisation assay for specific Mps1 activity. (A) The overall scheme of the FP assay; the peptide (green bar) is fluorescently labelled (yellow star) (B) FP measurements of WT Mps1 kinase activity (blue) by titrating the Bub1/Bub3 complex. Grey and black circles represent the measurement in the presence of NMS-P715 and Cpd-5, respectively.

Having established the conditions for the FP assay, we measured the ability of Cpd-5 and NMSP715 to inhibit the activity of Mps1 kinase domain on the KNL1 substrate, by titrating them into our assay mixture (Figure 2 and Table 2). From the titration curves, the IC_50_ values were estimated to be 9.2±1.6 nM for Cpd-5 and 139±16 nM for NMS-P715. Cpd-5 and NMS-P715 inhibit the C604Y variant with a significantly worse IC_50_ of 170±30 nM for Cpd-5 and 3016±534 nM for NMS-P715, respectively. A more moderate reduction was observed for the C604W substitution, with IC_50_ of 19±1 nM for Cpd-5 and 900±555 nM for NMS-P715. To validate the result of our new assay, the potency of the inhibitors was assessed also by probing phospo-KNL1 protein with a phospho-specific antibody for the C604Y mutant, which results in similar IC_50_ values (Supplementary figure 1A and Table 2). These results are fully in agreement with previous studies which showed that expression of the C604Y mutant renders cells more resistance to NMS-P715 than to Cpd-5 [11].

**Table 2.** Inhibition constants and affinities.

| IC<sub>50</sub> (nM) – cells |  |  |  |  |
| --- | --- | --- | --- | --- |
| Inhibitors |  | WT | C604Y |  |
| Cpd-5 |  | 17 | 320 |  |
| NMS-P715 |  | 170 | >2000 |  |
| IC<sub>50</sub> (nM) – αKNL1 |  |  |  |  |
| Inhibitors |  | WT | C604Y |  |
| Cpd-5 |  | 69 ± 23 | 458 ± 167 |  |
| NMS-P715 |  | 34 ± 6 | 1337 ± 402 |  |
| IC<sub>50</sub> (nM) – Bub1/Bub3 KNL1 FP assay |  |  |  |  |
| Inhibitors |  | WT | C604Y | C604W |
| Cpd-5 |  | 9.2 ± 1.6 | 170 ± 30 | 19 ± 1 |
| NMS-P715 |  | 139 ± 16 | 3016 ± 534 | 900 ± 55 |
| K<sub>D</sub> (nM) and ΔG<sub>calc</sub> (kJ/mol) – MST |  |  |  |  |
| Inhibitors |  | WT | C604Y | C604W |
| Cpd-5 | K<sub>D</sub> | 1.6 ± 0.2 | 471 ± 50 | 349 ± 81 |
|  | ΔG | 52.2 | 37.5 | 38.3 |
| NMS-P715 | K<sub>D</sub> | 4.7 ± 2.5 | 1764 ± 204 | 630 ± 115 |
|  | ΔG | 49.4 | 34.1 | 36.8 |
| Reversine | K<sub>D</sub> | 41 ± 21 | 86 ± 24 |  |
|  | ΔG | 43.8 | 41.9 |  |

**Figure 2:**
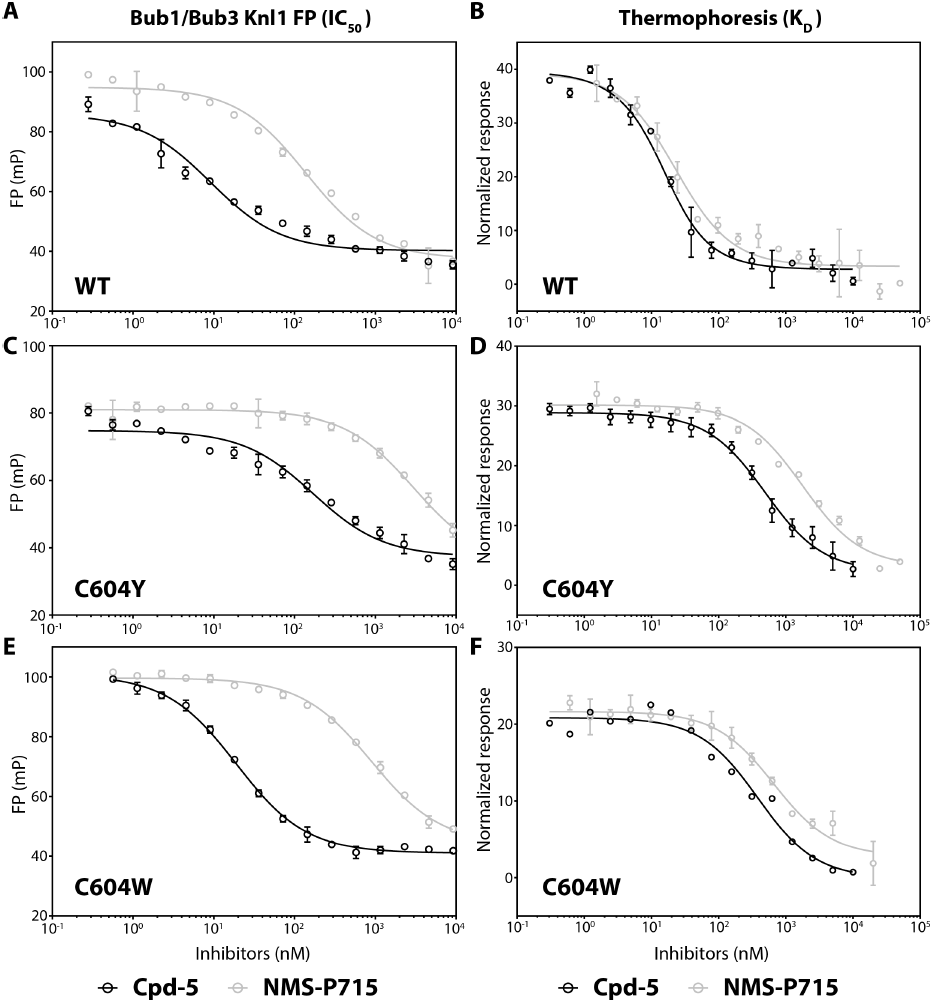
characterisation of Mps1 C604Y mutant and inhibitors. (A, C, E) FP assay for the Mps1 kinase variants in the presence of inhibitors; WT and C604Y/W kinase activities are shown in circles and triangles, respectively. IC_50_ values were calculated by titrating Cpd-5 (black) and NMS-P715 (grey). (B, D, F) Binding affinity measurements of the Mps1 variants with inhibitors, measured by MST.

### Binding affinities of Cpd-5 and NMS-P715 to WT and C604Y/W Mps1

The IC_50_ value shows how effective a compound is on a specific assay, but it is not a measurement of the binding of the inhibitor; the latter information is important to understand from a mechanistic perspective the action of inhibitors and the differential effect they assert to mutant proteins. Thus, to evaluate the influence of the C604Y/W substitution in the inhibition mode, the binding affinities were determined by microscale thermophoresis (MST; Figure 2 and Table 2). The MST results show that Cpd-5 binds approximately 200–300 fold better to the wild-type Mps1 (1.6±0.2 nM) than to the C604Y (471±50 nM) and C604W variants (349±81 nM); a similar trend is observed for NMS-P715, which binds to the wild-type Mps1 (4.7±2.5 nM), to the C604Y (1764±204 nM) and to the C604W variants (630±115 nM). Notably however, despite the significant reduction in binding affinity to the C604Y/s mutants, Cpd-5 binds better than NMS-P715. This observation is consistent with the IC_50_ measurements *in vitro* and with cell-based assays [11], where Cpd-5 shows 5-20 fold higher potency than NMS-P715 towards inhibiting the C604Y/W mutants. This is of potential interest for the discovery of compounds that target resistant variants of the kinase, and is important in light of our crystallographic structure models of Cpd-5 and of NMS-P715 bound to the C604Y/W mutants.

### Crystal structures of Mps1 kinase domain mutants bound to Cpd-5 and NMS-P715

The Mps1 C604Y/W kinase domain mutants C604Y and C604W were each co-crystalized with Cpd-5 and with NMS-P715. The crystal structures of the Mps1 C604Y variant in complex with Cpd-5 (PDB ID: 5MRB), NMS-P715 (PDB ID: 5NTT) and the Mps1 C604W with Cpd-5 (PDB ID: 5O91), were determined by molecular replacement, at 2.20 Å, 2.75 Å, 3.20 Å resolution, respectively, while the NMS-P715 bound to the Mps1 C604W structure (PDB ID: 5AP7) was reported in a previous study [12]. In all cases, there was clear electron density for Cpd-5 and NMS-P715 in the ATP-binding pocket (Figure 3). The overall structure of the protein adopts to a conformation very similar to the previously reported structures [2]. As in many structures of Mps1, the activation loop encompassing residues 676–685 had poor density, which only for the NMS-P715 was sufficiently well resolved to make a coarse model of the loop. Thr686 in the P+1 loop has been shown to be auto-phosphorylated in the recombinant protein [2]; although we tried to model the phosphoryl group of Thr686, it is not well resolved in the electron density map and therefore it has not been included in the structural models. Interestingly, the electron density indicated that a polyethylene glycol molecule (PEG), which sometimes encapsulates the catalytic Lys553 [2], is not present in our data. Instead, the side chain of Lys553 forms a hydrogen bond to the O atom adjacent to the diethyl phenyl ring of Cpd-5 and NMS-P715. This shows that ligand binding to Mps1 can be misrepresented if abundant PEG is used during crystallisation [36]. Both inhibitors are stabilized by two additional hydrogen bond interactions with the amide backbone of the “hinge loop” residue, Gly605. The electron density of the Cys604 point mutation is clearly resolved, indicating that the side chains of the substituted Tyr and Trp are well ordered in place.

**Figure 3:**
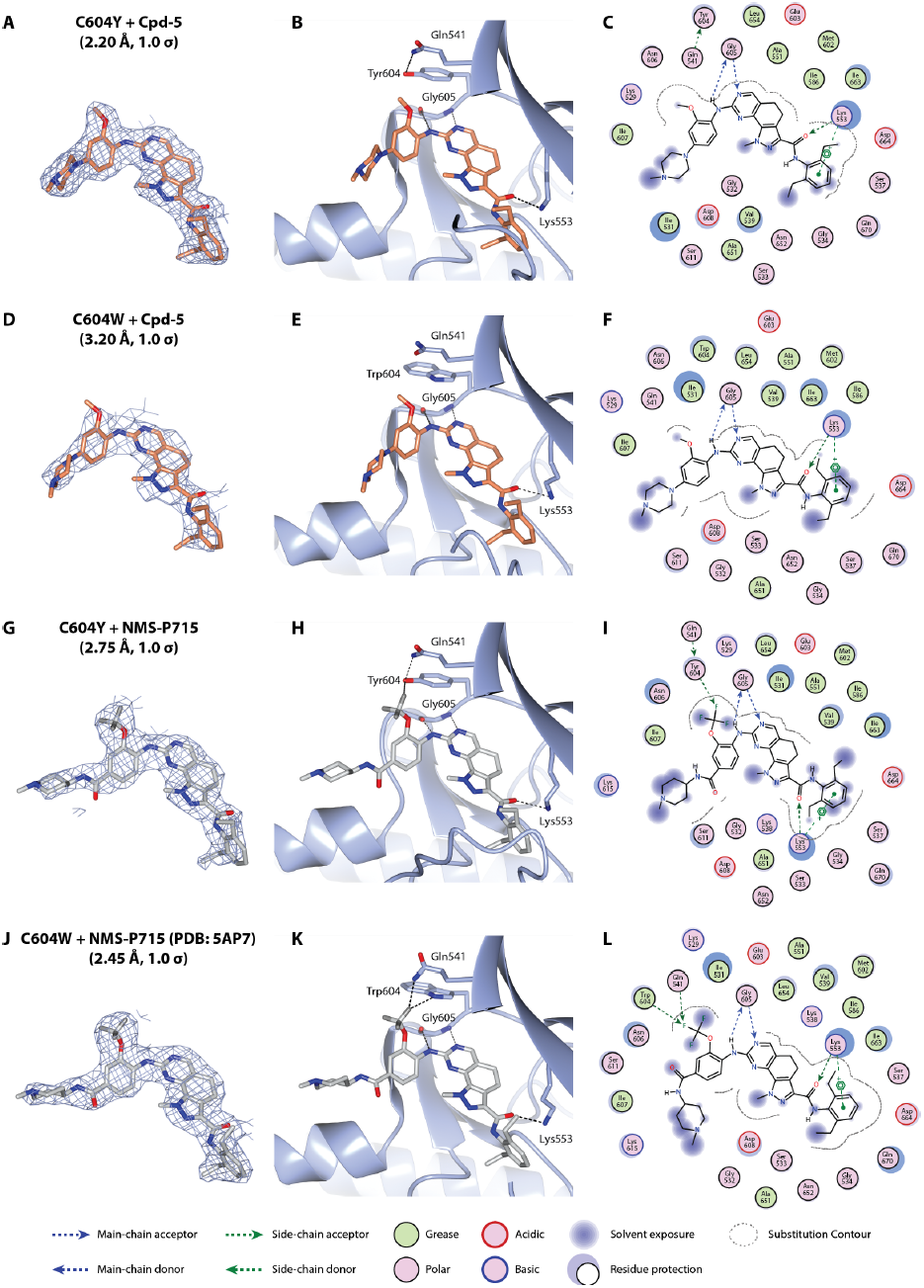
Comparison of the Cpd5 and NMS-P715 bound to C604Y/Wstructures. (A) *2mF*_*o*_*-DF*_*c*_ electron density map of Cpd-5 in Mps1 kinase contoured at 1.0σ (carved at 2.5 Å from the Cpd-5 atoms). (B) Cpd-5 in the ATP binding pocket of Mps1 C604Y; Cpd-5, the side chains of Gln541, Tyr604, and Lys553 as well as residue Gly605 are depicted as sticks; hydrogen bonds are depicted as black dotted lines. Figures were made in CCP4mg [38]. (C) Cpd-5 interactions with Mps1 kinase drawn by the Lidia routine in COOT. (D, E, F) The same as A, B, C for the interaction of Cpd-5 with Mps1 C604W. (G, H, I) The same as A, B, C for the interaction of Cpd-5 with Mps1 C604W. (J, K, L) The same as A, B, C for the interaction of NMS-P715 with Mps1 C604W

### Comparing crystal structures explains inhibitor resistance and selectivity

These structures, allow comparing the differences in the binding of different compounds to the Mps1 kinase domain (Figure 4). The key difference between Cpd-5 and NMS-P715 is the functional group attached on the phenyl moiety (Figure 4A, B). The trifluoromethoxy group of NMS-P715 and the smaller methoxy group of Cpd-5, are both positioned at close proximity to the side chains of Tyr/Trp604 and Ile531. The absence of the three fluorine atoms in the functional group of Cpd-5 lowers steric hindrance, and results in shifting the position of Cpd-5 towards Tyr/Trp604. The significantly lower affinity of NMS-P715 (2–3 fold lower in K_D_, corresponding to a 2–3 kJ/mol difference in ΔG_calc_, Table 2) can be attributed to the increased chances of steric hindrance between Tyr/Trp604 and Ile531 and the trifluoromethoxy moiety of NMS-P715.

**Figure 4:**
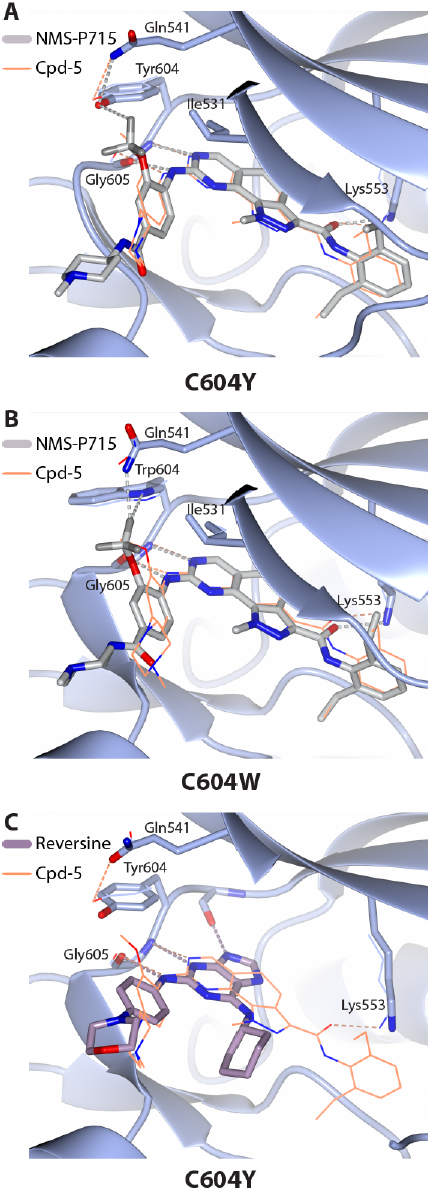
Comparison of the binding modes of Cpd5, NMS-P715 and reversine. (A) NMS-P715 and Cpd5 in the ATP binding pocket of Mps1 C604Y; (B) NMS-P715 and Cpd5 in the ATP binding pocket of Mps1 C604W; (C) Reversine and Cpd5 in the ATP binding pocket of Mps1 C604Y.

Structure comparisons also explain the molecular basis of the non-resistant phenotype of the C604Y mutant towards reversine [11] (Figure 4C and Table 2); the affinity of reversine to WT and the C604Y Mps1 variant is the same within error (Table 2 and Supplementary figure 1B). Comparison of the structure of the C604Y mutant bound to reversine (PDB: 5LJJ, [23]) clearly shows that there is no steric hindrance with the Tyr604 side chain, explaining why C604Y, and by deduction C604W, are non-resistant to reversine as previously reported [11,12].

We then compared the binding mode of the NMS-P715 and the Cpd-5 inhibitor to the Mps1 C604Y and C604W mutants (Figure 5). NMS-P715 binds better to the Trp mutant, than to the Tyr mutant. For NMS-P715 we observe that one of the fluorines when in complex with the C604W mutant makes a close contact with the Hε_1_ of Trp604 (2.3Å) and an even closer contact to the Hε_21_ of Gln541 (2.0Å). Particularly the latter close contact can be regarded as a good example of rare hydrogen bond of the C-F**⋯**H-N type [37]. The χ3-torsion angle of Gln541 changes by approximately 30 degrees to accommodate this interaction, in comparison to the other three structures. In the Tyr604 mutant however, these interactions are not formed; instead, a single hydrogen bond between Hε_21_ of Gln541 and the side-chain oxygen of Tyr604 is available. This explain why NMS-P715 binds better C604W; the calculated energy difference of ~3 kJ/mol based on the measured K_D_ (Table 1), is compatible with the energy of a weak hydrogen bond. Cpd-5 binds with similar affinity to both mutants; that is consistent with the lack of major differences in the binding mode of this inhibitor to both mutants.

**Figure 5:**
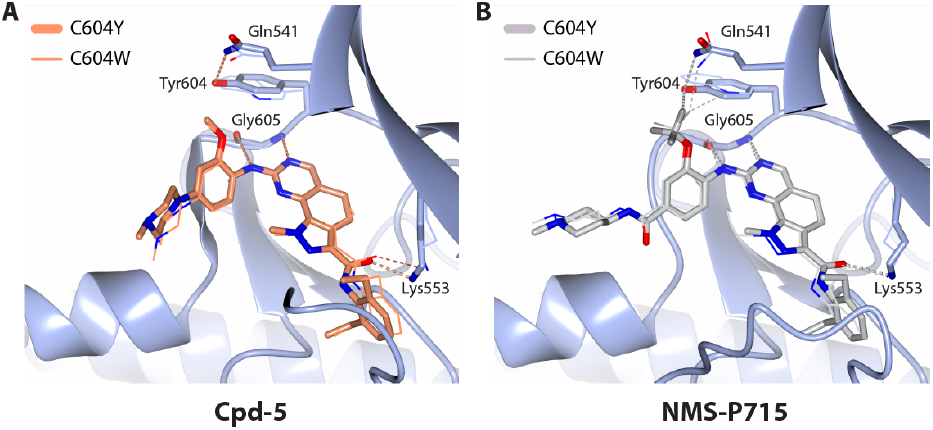
Comparison of the Cpd5 and NMS--P715 C604Y/W binding modes. (A) Cpd-5 in the ATP binding pocket of Mps1 C604Y/W; the C604W structure is shown with thin lines for comparison. (B) NMS-P715 in the ATP binding pocket of Mps1 C604Y/W.

### Conclusions and perspective

Our finding that binding of NMS-P715 is more affected than binding of Cpd-5 is consistent with cell studies showing that the C604Y substitution confers resistance more moderately to Cpd-5 than to NMS-P715 [11]. This also indicates that combinations of different Mps1 inhibitors can be used to avoid or combat resistance in the clinic, and molecular understanding of the Mps1 interaction with inhibitors is important. Based on the different favourable interactions in the methoxy group substitutions, we suggest that substitutions of the methoxy group could be used to develop, perhaps less potent, Mps1 inhibitors that could be used to target the Cys604 mutation, if the need arises in the clinic. For instance, inhibitors for the Tyr604 mutant may benefit from a single or a double fluorine substitution on the methoxy group rather than the triple substitution in NMS-P715. Concluding, this study contributes in understanding the mechanism of resistance in Mps1 kinase inhibitors, suggests a new efficient and specific assay to aid Mps1 inhibitor discovery, and puts forward novel design principles for the further development of this class of inhibitors.

## Acknowledgements

We thank Prof. Dr. Geert Kops for the many discussions about Mps1 kinase and his support. We thank the ESRF for their support, in particular beamline scientist David von Stetten, at beamline ID30A-3 and the MASSIF scientists for remote automated data collection.

## Author contribution statement

YH performed or participated in all experiments and made all figures; AK performed the antibody-based Mps1 activity experiments; NH helped in establishing and performing the FP experiments; FT helped in purification and crystallisation of the C604W mutant; RHM supervised experiments in cells; RPJ helped in structure refinement and analysis; AP analysed the structures and supervised this work; this work was initiated by AK and YH; YH and AP wrote the paper with help from all authors.

---

**figure S1:**
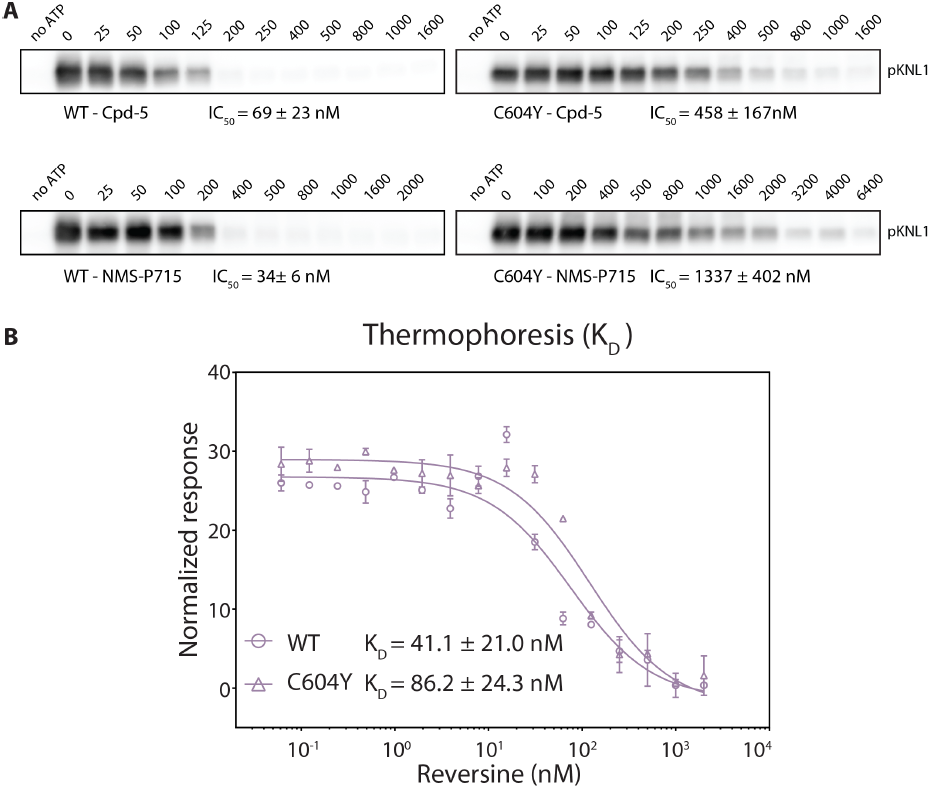
Mps1 kinase activity assays (A) IC_50_ measurements of the Mps1 kinase by probing phosphorylated KNL1 with antibodies. (B) Binding affinity measurements of the Mps1 variants with Reversine, measured by MST.

